# The evolution and spread of sulfur-cycling enzymes reflect the redox state of the early Earth

**DOI:** 10.1101/2022.08.05.502933

**Authors:** Katherine Mateos, Garrett Chappell, Aya Klos, Bryan Le, Joanne Boden, Eva Stüeken, Rika Anderson

## Abstract

The biogeochemical sulfur cycle plays a central role in fueling microbial metabolisms, regulating the Earth’s redox state, and impacting climate. However, geochemical reconstructions of the ancient sulfur cycle are confounded by ambiguous isotopic signals. Here, we use phylogenetic reconciliation to ascertain the timing of ancient sulfur cycling gene events across the tree of life. Our results suggest that metabolisms using sulfide oxidation emerged in the Archean, but those involving thiosulfate emerged only after the Great Oxidation Event. Our data reveal that observed geochemical signatures resulted not from the expansion of a single type of organism, but were instead associated with genomic innovation across the biosphere. Moreover, our results provide the first indication of organic sulfur cycling from the mid-Proterozoic onwards, with implications for climate regulation and atmospheric biosignatures. Overall, our results provide insights into how the biological sulfur cycle evolved in tandem with the redox state of the early Earth.

**Teaser:** Phylogenomics analyses reveal that the evolution of microbial sulfur metabolisms co-evolved with the redox state of the early Earth.

## Introduction

The biogeochemical sulfur cycle has played a crucial role in the evolution of life and surface processes over geologic time. Dissimilatory metabolisms, including elemental sulfur reduction, sulfate reduction, sulfate disproportionation, and sulfide oxidation fuel diverse microbes and play an important role in regulating the redox state of the surface of the Earth (*1*) (**Figure 1**). For example, the burial of biogenic sulfide in marine sediments may have contributed to progressive oxygenation of surface environments (*2*). Additionally, Earth’s carbon and sulfur cycles are linked through the metabolic reduction of sulfate, which is coupled with the oxidation of organic carbon. This process accounts for up to 50% of organic carbon mineralization (*3*). The sulfur cycle is also intricately entwined with cycling of other important elements, including nitrogen and various transition metals (*4, 5*). In particular, freely dissolved sulfide in seawater, especially during the Proterozoic eon (*6*), may have impacted the solubility of essential micronutrients such as molybdenum (*7*). Thus, a deeper understanding of the evolution of the biological sulfur cycle can offer important insights into the oxidation state of our planet over time and the evolution of other biogeochemical cycles.

**Figure 1.**
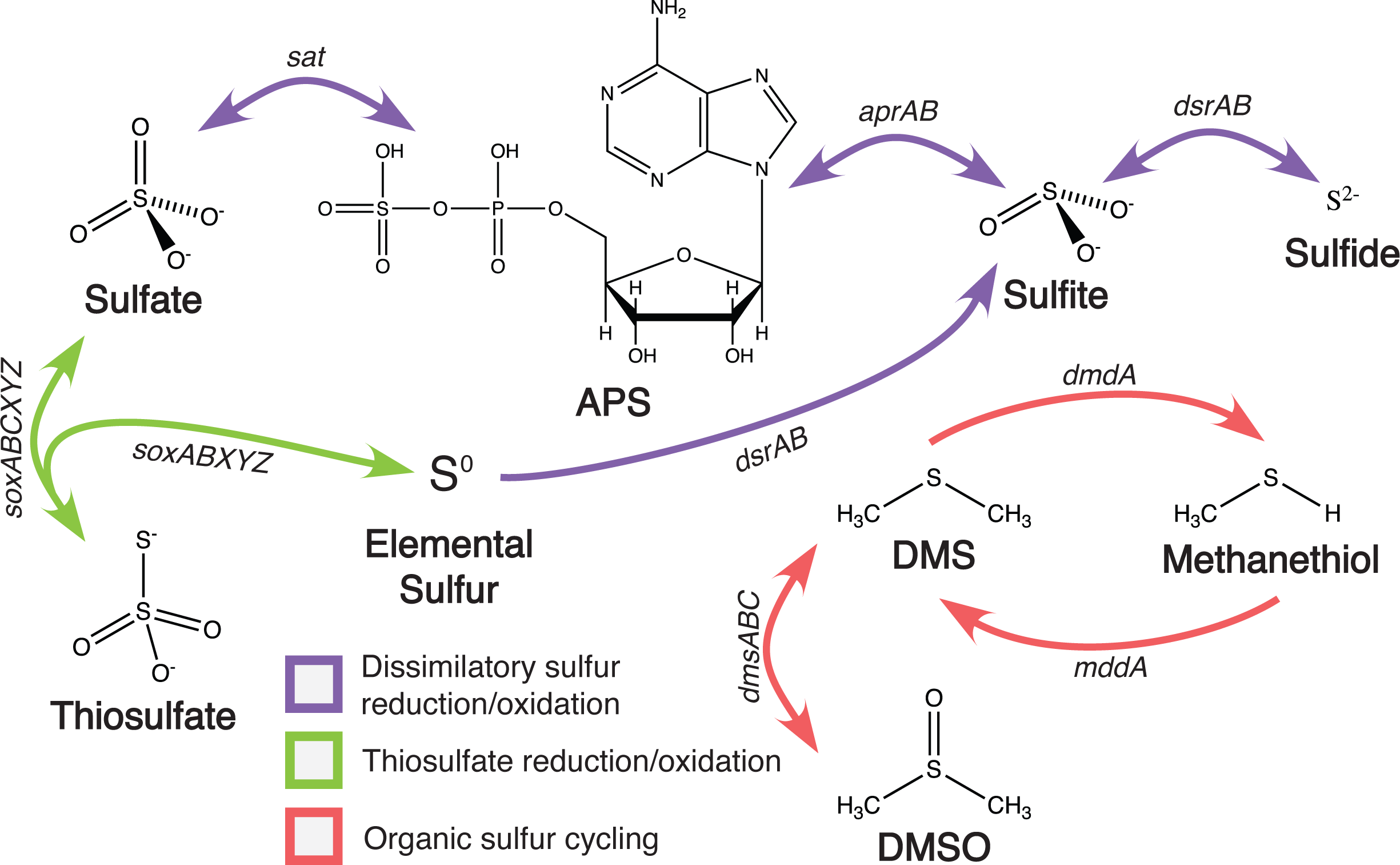
Schematic of the biological sulfur cycle, highlighting the genes included in this analysis (note that *sat* was excluded from the analysis because it is also used in several other pathways).

Several geochemical studies suggest that sulfur metabolisms were probably among the earliest microbial metabolisms on the ancient Earth. Early analyses of sulfur isotopes of pyrite and barite in the 3.5 billion year old (Ga) Dresser formation provided evidence for sulfate reduction in the Archaean era (*8*). While data from Philippot et al. (*9*) later argued that the isotopic fingerprint of the sulfides was more consistent with sulfur disproportionation, subsequent work supported the original claim (*10*). Since then, several studies have documented isotopic evidence of microbial sulfur cycling in a variety of Archean environments (e.g. (*11–13*)).

One of the biggest questions regarding the evolution of the biological sulfur cycle is how it co-evolved with the oxygenation of the Earth over time. It is now well established that the Earth’s surface underwent a major transformation at around 2.4 Ga, when atmospheric O_2_ levels increased above a threshold of 10^-5^ times present levels, known as the Great Oxidation Event (GOE) (*14*). As a consequence, volcanism was replaced by oxidative weathering as the major source of sulfur to the ocean, and this new sulfur source was predominantly in the form of sulfate, as opposed to volcanogenic SO_2_, dissolved sulfite, and the photochemical products S_8_ and sulfate (*15*). Further, O_2_ became abundant in surface waters as a potent oxidant of reduced sulfur species (*16*). The Archean-Proterozoic transition also witnessed a decline in hydrothermal activity on the ocean floor (*17*). Lastly, the deep ocean became fully oxygenated in the Neoproterozoic or early Paleozoic with the second rise of oxygen (*6, 18*), leading to enhanced sulfide oxidation within sediments (*19*). Geochemical data show isotopic expressions of these events in the sulfur cycle (*1*); however, it is so far unknown if these isotopic signals reflect merely an enhancement of a pre-existing process or true evolutionary innovations. This question has important implications for cause-effect relationships in Earth system evolution.

While most studies of the early sulfur cycle rely on geochemical analyses, this approach has limitations for three reasons: First, the isotopic signatures of differing sulfur metabolisms are not necessarily distinct enough to be recognizable in the sedimentary rock record. For example, analysis from the 3.2 Ga Moodies Group in South Africa confirmed the presence of reductive sulfur cycling, and hinted at the presence of oxidative sulfur cycling (*20*); however, the latter could not be unambiguously inferred. Second, the concentration of sulfate in the Archean ocean was low, possibly as low as 2.5 µM (*21*), dampening the signal of microbial sulfur isotope fractionation in the rock record. This challenge was illustrated by analysis of sulfur isotope ratios in a modern sulfate-poor analog of the Archaean ocean, where biological fractionations are muted despite the presence of active microbial sulfate reduction (*21*). Third, the Archean sulfur isotope record is famously impacted by photochemical processes acting on volcanogenic SO_2_ gas in an anoxic atmosphere (*22*). These photochemical reactions are recognizable by so-called mass-independent isotopic fractionations; however, distinguishing these from biogenic isotope effects requires analyses of all four stable isotopes of sulfur and relatively large sample sets (*10, 23, 24*). Thus, while the geochemical record has produced valuable insight into sulfur cycling in the Archean, the results can be inconclusive and often cannot distinguish between specific metabolic pathways.

Given the limitations and uncertainties presented by the geochemical record, pairing geochemical analysis with top-down phylogenetics approaches can provide important insights into the emergence and spread of distinct sulfur-cycling microbial metabolisms on the early Earth. Though no dedicated molecular clock work to date has been conducted on sulfur metabolisms, genomics-based studies suggest that dissimilatory sulfite and sulfate reduction and sulfide oxidation emerged as early forms of energy metabolism (*25–28*). Phylogenetics analyses suggest that the *dsr* genes, which catalyze the reduction of sulfite to sulfide or in the reverse for sulfide oxidation, were among the first sulfur-based genes to arise (*25, 27*), with the reductive form arising first (*29*). Anaerobic anoxygenic photosynthesizers would likely have emerged soon thereafter, possibly using these genes to oxidize sulfur and sulfide (*30, 31*). Genomics analyses also indicate that sulfur disproportionation likely arose subsequently (*27, 28*). As the oceans became increasingly oxygenated, thiosulfate would have become more available, paving the way for the evolution of thiosulfate oxidation/reduction via the *sox* pathway (*26*).

However, while phylogenetics-based approaches generally track the birth of specific lineages or genes, the birth of a gene does not necessarily coincide with the time at which the function of that gene became ecologically important. As genes are horizontally transferred between divergent microbial lineages, the genes that serve a useful function are the ones most likely to be retained in a genome (*32–37*). Thus, an increase in horizontal gene transfer (HGT) events for a particular gene at a specific point in time likely gives an indication that the gene in question was ecologically important during that time period. Similarly, speciation events indicate that the lineage encoding a specific functional gene, and therefore the metabolism it facilitates, has expanded into new ecological niches.

To examine the evolution of the biological sulfur cycle over time, we therefore used phylogenomics approaches to track the timing of speciation, duplication, loss, and horizontal gene transfer events for sulfur cycling genes across a time-calibrated tree of life. This analysis allows us to determine approximately when these genes first arose and then proliferated across the tree of life on the early Earth. A similar analysis of nitrogen-cycling genes revealed that nitrogen fixation arose and spread early, while genes related to denitrification from nitrite arose and spread much later in Earth history (*38*). Here, we focus on constraining the timing of speciation, duplication, loss, and horizontal gene transfer events for genes related to dissimilatory sulfate reduction and sulfide oxidation via sulfide, transformations between sulfate and thiosulfate, as well as organic sulfur cycling.

## Results

### Construction of species tree and time-calibrated chronogram

We constructed a species tree from an alignment of fifteen universal single-copy ribosomal genes in order to conduct the phylogenetic reconciliation (**Supplementary Figure 1**). The resulting tree placed the Eukaryotes within the Archaeal domain, consistent with a two-domain tree of life, as has been recovered previously using similar methods (*39–41*). We constructed chronograms from this species tree using two autocorrelated clock models (lognormal (LN) and Cox-Ingersoll-Ross (CIR)) and one uncorrelated gamma multiplier (UGAM) clock model. We tested both liberal and conservative fossil calibration points to construct the molecular clock as a sensitivity test (**Table 1**; see Methods). For all clock models, the liberal calibration points returned unrealistic ages (i.e. between 5.3-6 Ga, prior to the formation of the Earth) for the last universal common ancestor (LUCA) (see chronograms with error bars shared FigShare at figshare.com/projects/The_evolution_and_spread_of_sulfur-cycling_enzymes_across_the_tree_of_life_through_deep_time/144267). Thus, we only used the conservative calibration points for the remainder of this analysis. Using the conservative calibration points, the LN clock returned a LUCA age of approximately 4.48 Ga, the CIR clock returned a LUCA age of approximately 4.05 Ga; and the UGAM clock returned a LUCA age of approximately 3.93 Ga. We report the results from the CIR clock in the main text because autocorrelated clock models have previously been shown to outperform uncorrelated models such as UGAM (*42, 43*); among the autocorrelated models, the CIR results were closer to most reports for the approximate age of LUCA (approximately 3.8 Ga (*44*)) than the results from the LN clock model. All results from the LN and UGAM clocks are reported in the Supplementary Materials. All Newick files, alignments, and chronograms with error bars have been deposited in FigShare at https://figshare.com/projects/The_evolution_and_spread_of_sulfur-cycling_enzymes_across_the_tree_of_life_through_deep_time/144267.

**Table 1.**
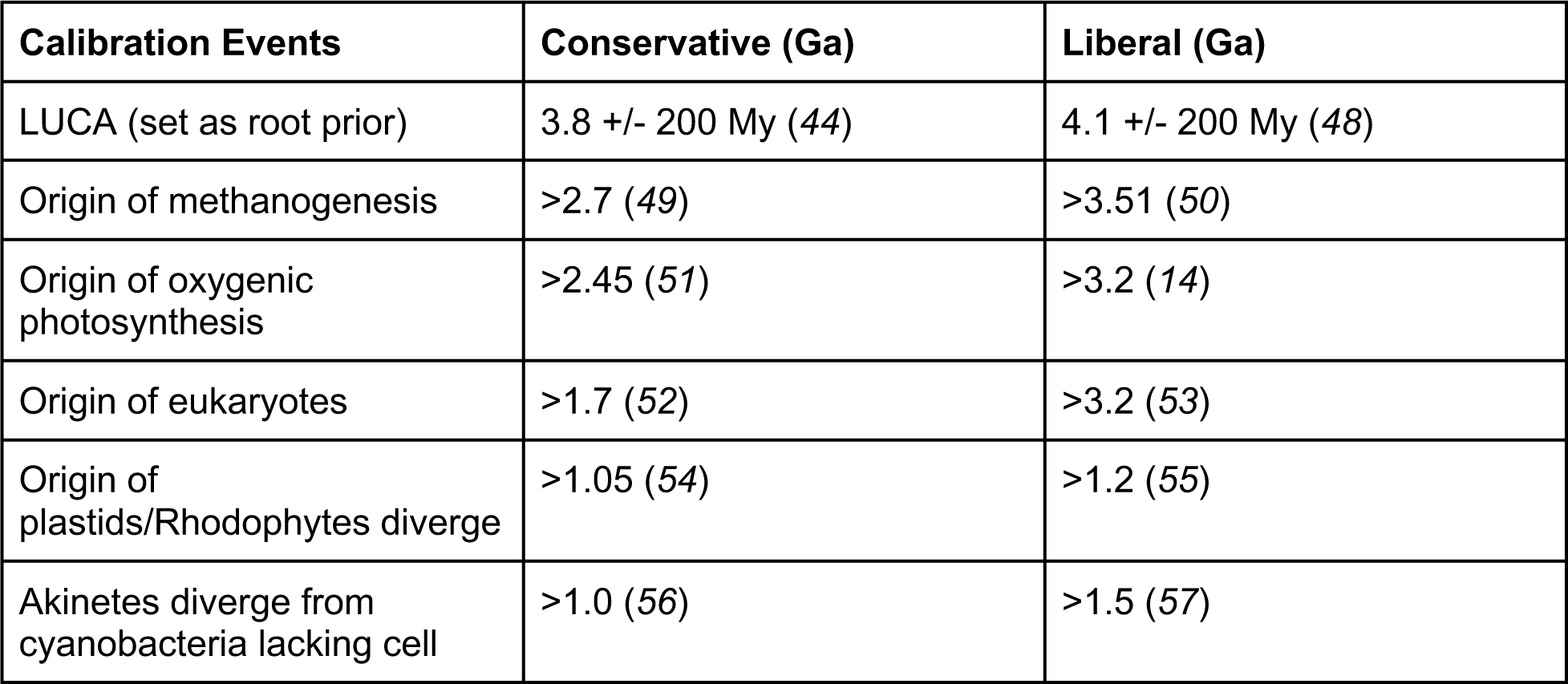
Fossil calibration points used for calibrating molecular clocks. Calibration points were set as the hard constraint in Phylobayes, indicating the latest date by which a specific clade must have split. The “conservative” time points reflect the dates for which there is the most consensus; “liberal” time points reflect the earliest date reported in the literature. Note that the root prior (LUCA) was set using a gamma distribution with mean 3.8 Ga (conservative, (*44*)) or 4.1 Ga (liberal, (*48*)) and standard deviation of 200 My.

### Phylogenetic distribution of sulfur-cycling genes

We used AnnoTree (*45*) to determine the distribution of sulfur-cycling genes across the tree of life. Genes related to dissimilatory sulfate reduction and sulfide oxidation via sulfite, including *dsr* and *apr* genes, tended to be fairly widespread across the tree of life: *aprAB* in particular is fairly widespread, occurring in approximately 47 bacterial and 5-6 archaeal phyla, and *dsrAB* is found in 32 bacterial and 4-5 archaeal phyla. In contrast, genes related to thiosulfate oxidation/reduction, particularly the *sox* group of genes, were more phylogenetically restricted, occurring in approximately 14-20 bacterial and 1 archaeal phylum, with the majority of gene hits restricted to the Proteobacteria superphylum. The exceptions to this rule were *soxB* and *soxC*, which were much more widespread across the tree of life, occurring in approximately 31-38 bacterial and 4 archaeal phyla. Finally, the organic sulfur cycling genes *dmdA*, *dmsA*, and *mddA* each displayed a different phylogenetic distribution: *dmdA* was found in only 13 bacterial and 2 archaeal phyla, restricted mostly to the Proteobacteria and Actinobacteria; *dmsA* was much more widespread, identified in 43 bacterial and 6 archaeal phyla, but generally not observed in the Patescibacteria; and *mddA* was similarly widespread, found in 34 bacterial and 4 archaeal phyla, most noticeably absent from the Patescibacteria and the Firmicutes phyla.

### Identification of duplication, loss, and horizontal gene transfer events for sulfur-cycling genes

For the 13 genes of interest, we quantified gene speciation, duplication, loss and horizontal gene transfer events using multiple reconciliation algorithms. Reconciliation was performed by comparing the topology of the maximum likelihood gene trees for each gene to fossil-calibrated chronograms using three different clock models (CIR, UGAM, and LN) using the reconciliation programs AnGST (*46*) and ecceTERA (*47*). The overall trends we observed were the same between AnGST and ecceTERA. The results we report here are those from ecceTERA, which can take into account both sampled and unsampled (including extinct) lineages (*47*). The analyses presented below focus on results from ecceTERA based on the CIR clock model, with replicate analyses with similar results from ecceTERA using the UGAM and LN clock models presented in **Supplementary Tables 1-2 and Supplementary Figures 2-6**.

In interpreting the dates for events identified through phylogenetic reconciliation, it is important to consider the limitations inherent in dating each of these gene events. The phylogenetic reconciliation identifies branches on which events occurred, and thus there is no way to determine when on a given branch a specific gene event occurred. This can be more clearly visualized in **Figure 2** and **Supplementary Figures 2-3**, which show the full branch lengths along which events could have occurred. Early events tended to be more well-constrained than late events partly because most early events were speciation events, which occur at specific nodes on the tree; additionally, earlier loss/duplication/HGT events occurred on shorter internal nodes rather than on the leaves of the branches, which occurred later. This was likely due to the taxonomic sampling included in the tree, in which one representative from each class was represented; a tree with a large number of more closely related taxa (i.e. at the species or subspecies level) would have shorter branches at the tips but would be computationally intractable to create. Combined with the inherent error associated with dating events on chronograms dating back billions of years, estimates of when gene events occurred should not be taken as absolute dates. Instead, we emphasize the relative timing of these events. By examining the distributions of when specific gene events occurred, we are able to better understand the relative timing of when specific metabolisms became ecologically important. The histograms presented in **Supplementary Figures 4-6** present the same data as **Figure 2** and **Supplementary Figures 2-3,** but instead depict the proportion of total gene events within each time bin according to the midpoint of the time range along which an event could have occurred. This depiction facilitates the visualization of trends. **Figure 3** depicts a simplified summary of the results, representing each gene event as a single point defined by the midpoint dates of the time range along which an event could have occurred. The earliest events for each gene are reported in **Table 3**.

**Figure 2.**
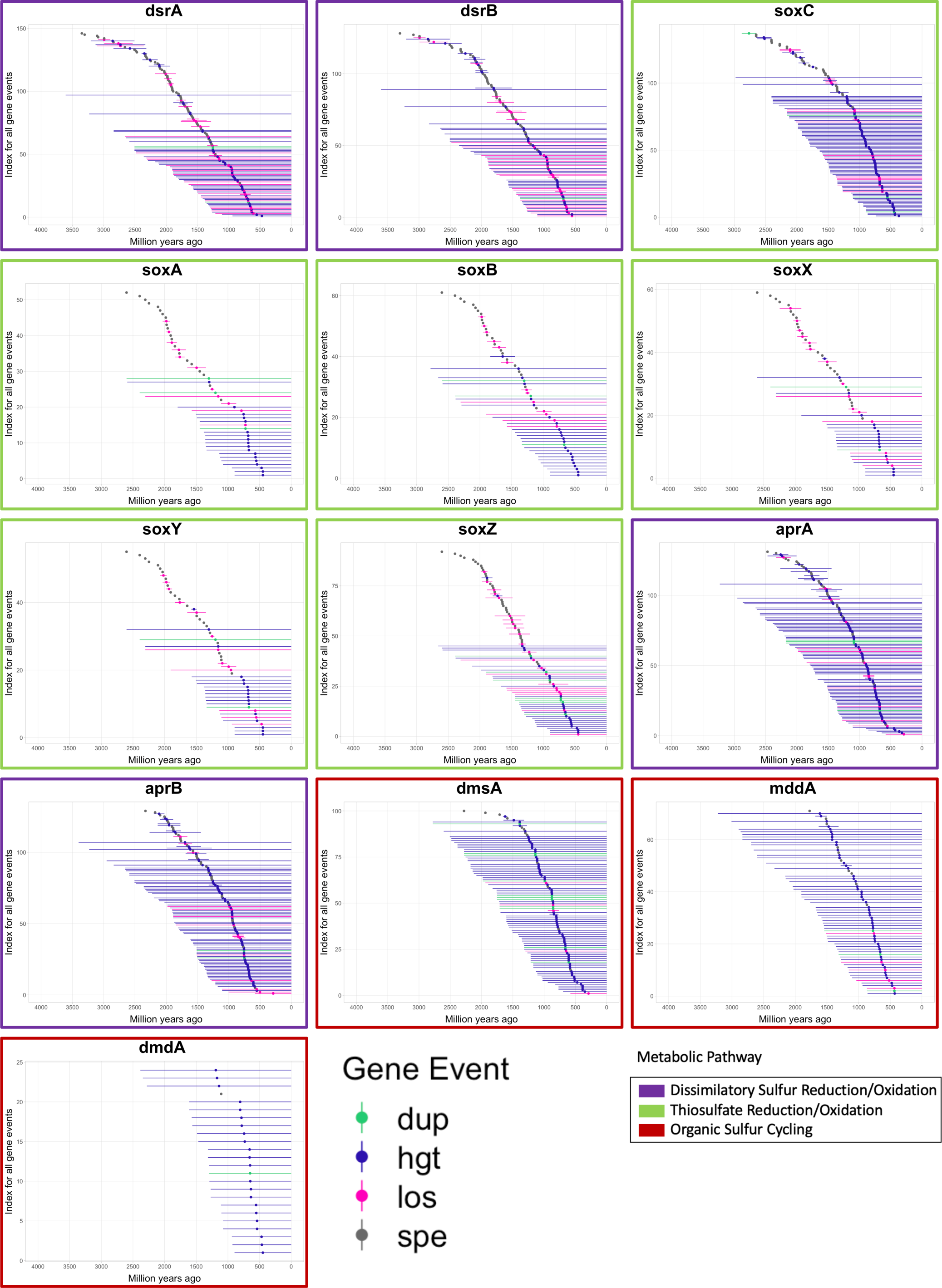
Time ranges for each gene duplication, transfer, loss, and speciation event identified for all sulfur-cycling genes. The time ranges reflect the branch lengths for branches on the time-calibrated tree of life on which these events were identified; the event could have occurred anywhere along the branch. Reconciliations were conducted with the CIR clock model. Graphs are placed in chronological order according to the midpoint of the earliest event. Points are colored according to the type of gene event. Colored boxes outlining each graph represent the general metabolic pathway in which each gene belongs. Dup =duplication, hgt = horizontal gene transfer, los = loss, spe = speciation event.

**Figure 3.**
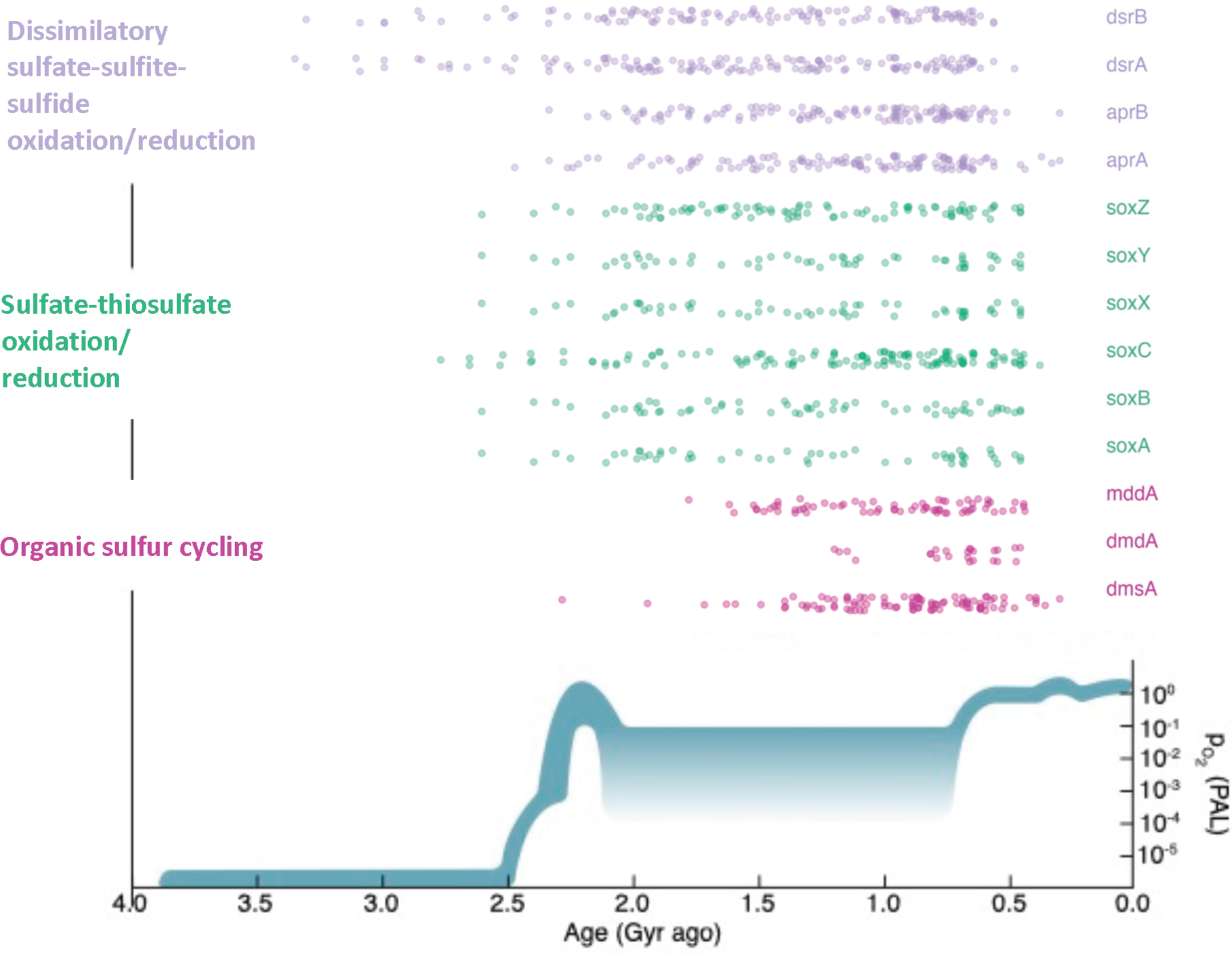
Dot plot showing the timing of gene loss, duplication, speciation, and HGT events for sulfur cycling genes as identified by ecceTERA using the conservative CIR clock model. Points are vertically jittered to facilitate visualization of individual events. A plot of changes in atmospheric partial pressure of oxygen over time was adapted from Lyons et al. (2014) for comparison purposes with the timing of the Great Oxidation Event (GOE).

The gene events are largely dominated by speciation events early in the evolution of each gene, often followed by a series of gene losses, then later dominated by horizontal gene transfers (**Figure 2; Supplementary Figures 2-6**). According to the earliest events identified for each of the genes in **Table 3** and as depicted in **Figures 2 and 3**, the most ancient genes include *dsrA* and *dsrB,* which catalyze the oxidation and reduction of sulfite and sulfide. (See **Figure 1** for a schematic of the sulfur cycle and the steps catalyzed by each of these genes.) The gene *apr*, which catalyzes the oxidation and reduction of adenylyl sulfate (APS) and sulfite, had more varied results depending on the clock model used, but according to the CIR clock model emerged later than *dsr* and around the same time as the *sox* genes. The *sox* genes, which are involved in sulfate and thiosulfate (S_2_O_32-_) oxidation, generally show a marked rise in speciation events beginning around 2.4 Ga, followed by a second rise in HGT events around 1.2 Ga (though this is less well-constrained in time) (**Figures 2 and 3**). This trend is fairly consistent across all *sox* genes, with the exception of *soxC*, which had many more gene hits and gene events than the rest of the *sox* genes, and was found in a wider range of organisms overall; reconciliation revealed an earlier emergence and proliferation compared to the other *sox* genes. It is important to note that reconciliation using the uncorrelated UGAM clock model showed that the *sox* genes had earlier gene events than *dsrB* and *aprAB* (**Supplementary Figures 3 and 4; Supplementary Table 2).** This was mostly driven by the speciation events; gene losses, transfers and duplications for the *sox* genes generally occurred later than for the *apr/dsr* genes.

For all clock models, the organic sulfur cycling genes appeared to be much younger than the genes involved in sulfur oxidation and reduction for energy metabolism. The gene *dmdA* converts dimethyl sulfide ((CH_3_)_2_S, DMS) into methanethiol (CH_3_SH), *dmsA* converts DMS to dimethyl sulfoxide ((CH_3_)_2_SO, DMSO), and *mddA* is involved in the conversion of methanethiol (CH_3_SH) to dimethylsulfide ((CH_3_)_2_S). While the organic sulfur cycling genes had varying timelines for their initial gene events, some as far back as 2.6 Ga, the number of gene events did not meaningfully begin to rise for any of these genes until approximately 1.6 Ga (**Figures 2 and 3**).

## Discussion

Our analysis of gene duplication, speciation, transfer, and loss events for sulfur cycling genes across Earth history provides insight into the relative timing for the proliferation of these genes across the tree of life, and thus has implications for when specific sulfur metabolisms became ecologically important. As has been suggested previously, if a gene is acquired via horizontal gene transfer and retained in the genome, this indicates that the horizontally transferred gene likely has been selected and retained because it performs a useful ecological function (*32–37, 60*). Thus, a rise in horizontal gene transfer events for a specific gene at a given time can indicate when these genes became favorable or ecologically useful given the conditions of the environment at that point in Earth’s history (*38*). Similarly, a speciation event occurs when a lineage containing that gene splits into two species, indicating that the metabolic pathway including that gene has begun to expand into new ecological niches and taxa.

It is important to note that many of these events occurred on long branches terminating in leaves of the species chronogram, meaning that these specific events could have occurred anywhere on that branch, up to the present day. Due to limitations inherent in gene-tree-species-tree reconciliation, the date of each event can only be confined to the branch of the species tree where it occurred, meaning that it could have happened at any point in time between the two nodes of that branch (as illustrated in **Figure 2** and **Supplementary Figures 2-3**). Caution is particularly warranted in interpreting the apparent peak in the number of events for many genes at ∼750 Ma as shown in **Figure 3** and **Supplementary Figures 4-6**. Many of these events occurred on leaves of the phylogenetic tree (as opposed to internal nodes), and the gene duplication, transfer or loss events occurred at some point between the terminus of the leaf and the last node of that leaf, many of which occurred at approximately 1.5 Ga. Thus, although the midpoints of these events were calculated to have occurred around 750 million years ago, in reality these events occurred at some point during a prolonged time period that extends to the present day, as shown in **Figure 2**. Due to these limitations in the data, caution is warranted in interpreting the timing of these gene events, as the calculated dates are rough estimates by necessity. Thus, our analysis emphasizes relative trends in the frequency of events over time, rather than focusing on precisely dated events. Additionally, the approximate dates for gene birth events are inferred based on the earliest event for that gene in the reconciliation, so the birth of that gene occurs prior to the earliest event by definition.

### Dissimilatory sulfate and sulfite reduction and sulfide oxidation

Dissimilatory sulfide oxidation involves the genes *dsrAB*, with sulfite as an intermediate. The same gene is also involved in sulfite reduction back to sulfide. According to the two autocorrelated (and thus more reliable (*42*)) clock models, these genes show their first events around ∼3.5 Ga (**Figures 2 and 3**). Sulfide would have been readily supplied by hydrothermal vent systems in the Archean ocean (*61*), and biological oxidation of sulfide may have been coupled to the reduction of Fe^3+^ or trace O_2_ that occurred locally in the surface ocean. The findings that the *dsrAB* genes have an ancient origin are consistent with phylogenetic results from Wagner et al. 1998, who used targeted gene sequencing and 16S rRNA sequencing techniques to show that dissimilatory sulfite reductase genes originated at around 3 Ga (*62*). Moreover, sulfite would have been supplied naturally by dissolution of volcanogenic SO_2_ in water (*15*). The reduction of sulfite could have been coupled to organic matter oxidation, as well as to volcanogenic or biogenic H_2_ oxidation.

Dates for the rise and spread of *aprAB*, which is involved in dissimilatory sulfate reduction with adenylyl sulfate (APS) as an intermediate, varied more widely across clock models and is therefore more difficult to constrain. The supply of sulfate to the Archean ocean, prior to the GOE, would have been limited to photochemical SO_2_ oxidation (*12, 22*). Nevertheless, geochemical data suggest that dissimilatory sulfate reduction is ancient, going back to 3.5 Ga (*8*) Previous studies have theorized that both the *apr* and *dsr* genes were involved in early oxidative pathways using sulfide in ancient microbial mats around 3 billion years ago (*26*).

While the oldest events for *dsr* date back to the Archean, our data also reveal an expansion for this gene around the GOE and again in the later Proterozoic or early Phanerozoic, concurrent with genes involved in dissimilatory sulfate reduction. The Paleoproterozoic events are likely linked to the increasing supplies of sulfate due to enhanced rates of oxidative weathering under an oxygenated atmosphere. The late-Proterozoic events may be linked to oxygenation of the deep ocean and associated growth of the marine sulfate reservoir (**Fig. 3**). Sulfate reduction and sulfide oxidation would have become more favorable metabolic pathways under these conditions (*63*). Our data thus suggest that these geochemical transformations of Earth’s surface directly impacted biological evolution.

### Sulfate-thiosulfate transformations

The Sox enzyme system is involved in the reduction and oxidation of sulfate and thiosulfate, respectively (*64*). A version of this pathway, omitting the SoxCD complex, can also be used to oxidize hydrogen sulfide to elemental sulfur (*1*). Most *sox* genes arose and began to speciate around or after the time of the GOE, approximately 2.4 Ga. Subsequently, the majority of the horizontal gene transfer events associated with the *sox* genes did not occur until much later, at approximately the time of the Neoproterozoic Oxygenation Event (NOE, (*14*)), in which the deep ocean is thought to have become more pervasively oxygenated approximately 850-540 million years ago (**Figure 3**). The rise in the number of speciation events for *soxABXYZ* approximately 2 billion years ago approximately coincides with increasing sulfate availability in the Earth’s oceans after the GOE (*65*). Thiosulfate has an intermediate redox state (S(+II)) between sulfide (S(-II)) and sulfate (S(+VI)) and forms most commonly during microbial sulfide oxidation (*66*). Hence the expansion of *sox* genes across the tree of life in the Paleoproterozoic is most parsimoniously attributed to increasing availability of O_2_ and therefore enhanced sulfide oxidation. This finding is consistent with geochemical evidence for enhanced disproportionation of elemental sulfur in the mid- to late-Proterozoic (*16, 19*), as elemental sulfur, like thiosulfate, is an intermediate in microbial sulfide oxidation. Moreover, the rise in horizontal gene transfer events around the time of the NOE, while not as well-constrained in time, suggest that increasing oxygen levels enabled the expansion of the biological sulfur cycle. In other words, these results indicate that increasing oxygen on Earth paved the way for the expansion of niche space and innovations in microbial evolution.

We identified many more events for the gene *soxC* compared to the other *sox* genes, many of which occurred earlier than the other *sox* genes. While it is unclear why this is the case, it may be related to the fact that *soxC* is not involved in the alternate *sox* pathway that creates elemental sulfur from sulfide. The *soxC* gene is part of a sulfur dehydrogenase molybdenum enzyme complex called *soxCD* that catalyzes a six-electron transfer in the middle of the *sox* sequence, and appears to be reliant on the other enzyme complexes in the *sox* sequence (*67, 68*). *soxC* exists in a wider range of organisms than the rest of the *sox* genes, which were primarily found in Proteobacteria. We speculate that this pattern may be the result of the gene’s relationship to another gene with a similar function, *sorA*, which has a 26.5% sequence identity to *soxC* (*68*) and is similarly widespread across the tree of life. Alternatively, it could indicate a separate function beyond the *sox* pathway for *soxC* that would require further investigation.

### Organic sulfur cycling

The organic sulfur cycle involves the biological formation of volatile organic compounds such as dimethyl sulfide (DMS) and methanethiol. The genes *dmdA* and *dmsA,* which are the key enzymes involved in DMS metabolisms, record their first events approximately 1.5-2 Ga, possibly linked to the rise of eukaryotic algae, whose production of organic sulfur gasses has been implicated in global cooling in the late Proterozoic (*69*). Similarly, the gene *mddA*, which converts methanethiol into DMS, also seems to have emerged and proliferated approximately 1.5-2 Ga. Thus, our results suggest that bacteria were capable of generating and metabolizing DMS only after the GOE, possibly with important implications for climate regulation on the early Earth, because DMS particles are known to act as cloud condensation nuclei, which has been hypothesized to cool the Earth’s surface (*70*). We speculate that DMS-generating metabolisms arose in response to a larger sulfate reservoir in the ocean from the GOE onwards, which may have led to organic-matter sulfurization in diagenetic settings (e.g., (*71*).

Moreover, volatile organic sulfur compounds such as DMS are important as potential remotely detectable biosignatures, because they could conceivably be detected on other planets with an anoxic biosphere using analysis of the spectral signatures of the planet’s atmosphere (*72*). Our results thus suggest that Earth’s biosphere may potentially have been detectable through this technique only within the past 1.5-2 Ga.

Shifts in the biological sulfur cycle have a profound impact on the global carbon cycle and Earth’s climate, and are closely tied to the redox state of the Earth. Our results suggest that microbial energy acquisition via sulfite reduction and possibly sulfide oxidation emerged early in Earth history, which is consistent with volcanic and hydrothermal sources of sulfite and sulfide, respectively. While our results cannot confirm geochemical evidence of microbial sulfate reduction (driven by *apr*) going back to 3.5 Ga, the dates obtained for this gene vary widely between clock models, and thus our results do not preclude the possibility that sulfate reduction arose earlier, and will require further investigation. We also find that metabolisms involving intermediates such as thiosulfate proliferated across the tree of life only after the Paleoproterozoic Great Oxidation Event, as the Earth’s ocean and atmosphere became more oxidizing. However, our analysis goes beyond the geochemical records, because our data reveal that the expressions of these geochemical signatures were not merely the result of preservation or expansion of a single organism but instead caused by the radiation of genomic innovations across the tree of life. We further show that the growth of the marine sulfate reservoir after the GOE triggered an expansion of organic sulfur metabolisms, which would have added an important biosignature to Earth’s atmosphere from the Proterozoic onwards. Lastly, our analyses reveal an expansion in all sulfur metabolisms around the Neoproterozoic, highlighting that this time period not only witnessed the rise of eukaryotic life but was also an important driver of microbial evolution.

## Materials and Methods

### Genome selection and construction of species tree

To construct the species tree, we included one representative genome from each bacterial and archaeal order, based on GTDB taxonomy (*73, 74*). Some eukaryotic genomes were also included in order to create a full tree of life in order to capture putative gene transfer events between the archaeal and bacterial domains, and to include additional time calibration points in the eukaryote domain for the molecular clock. However, the focus of the study was on sulfur cycling genes within bacterial and archaeal genomes. GToTree (*75*) was applied to identify and align single-copy universal ribosomal genes from the genomes we selected. The concatenated gene alignments were created from a set of sixteen universal single-copy genes (*76*), and we excluded genomes with fewer than half of the single-copy genes. Briefly, the GToTree workflow used prodigal (*77*) to predict genes on input genomes, then identified genes with HMMER3 v3.2.2 (*78*), individually aligned genes with MUSCLE v5.1 (*79*), trimmed the alignment with trimal v1.4.rev15 (*80*), and concatenated aligned genes with FastTree2 v2.1.1 (*81*). The resulting alignment was used to construct a phylogeny using RAxML v. 8.2.9 (*82*) with 100 rapid bootstraps using the PROTGAMMALG model of evolution as per (*76*). The root of the tree was placed in between the archaeal and bacterial domains by designating the entire bacterial domain as the outgroup. The resulting tree contains 871 genomes, including 777 bacterial, 80 archaeal, and 14 eukaryotic genomes.

### Construction of time-calibrated chronogram

The species tree was converted to a chronogram using Phylobayes v4.1b (*83*). We tested two separate sets of calibration points, one conservative (which represents the earliest date for which there is the most consensus for a given event based on the current scientific literature) and one liberal (which represents the earliest date for which there is any evidence of a given event based on the current scientific literature) to test the sensitivity of methodology (**Table 1**). The root age was set via a normally distributed gamma root prior according to dates specified in Table 1 with a standard deviation set to 200 million years, consistent with previous studies (*84*).

To generate chronograms, we tested three different clock models: autocorrelated log normal (LN) (*85*), uncorrelated gamma multiplier (UGAM) (*86*), and the autocorrelated Cox-Ingersoll-Ross (CIR) process (*42*). For each model and set of calibration points, two chains were run concurrently and were compared as a test of convergence. We analyzed convergence using the tracecomp program in Phylobayes, requiring an effective size >100 and a maximum difference between chains of <0.3. Each chain was run for >60,000 cycles. Chronograms were generated using the readdiv function in Phylobayes, discarding the first 2500 cycles as a burn in.

### Identification of sulfur cycling genes and construction of gene trees

We identified sulfur cycling genes of interest using the sulfur metabolism pathway on the Kyoto Encyclopedia of Genes and Genomes (KEGG) (*87–89*). In line with previous genomic studies of the sulfur cycle, we analyzed dissimilatory sulfate reduction/sulfide oxidation genes (*aprAB*, *dsrAB*) (Anantharaman et al. 2018) and thiosulfate oxidizing/sulfate reducing genes (*soxABCXYZ*) (Canfield et al. 2010). Though the *sat* gene catalyzes the first step of dissimilatory sulfur cycling, we excluded this gene from our study because it is also used in other metabolic pathways that were not of specific interest here. We also examined genes that were involved in the production of volatile organic sulfur compounds, including methanethiol, dimethyl sulfoxide and dimethyl sulfide (*mddA, dmdA, dmsA*). The list of sulfur-cycling genes analyzed here was not meant to be exhaustive, but rather focused on core genes involved in dissimilatory sulfur oxidation and reduction for energy acquisition as well as select genes involved in organic sulfur cycling.

For consistency in identifying genes across genomes, we used Annotree (*45*) to identify sulfur cycling genes in microbial genomes using KEGG orthology numbers as queries (**Table 2**). We limited our analysis to the core dissimilatory sulfur cycling and thiosulfate reduction and oxidation genes that were included in the Annotree database. It is important to note that some metabolic pathways use the same genes for catalyzing oxidation and reduction reactions, and thus cannot be distinguished using these methods. For example, sulfur disproportionation uses the same enzymatic pathways as sulfate reduction (*28, 90*). The default Annotree settings (minimum 30% identity, maximum e-value of 10^-5^, minimum 70% subject and query alignment) were applied for identifying genes in genomes. Annotree output was curated to only include genes from genomes within our species tree. Note that although eukaryotic genomes were included in the species tree, AnnoTree does not include eukaryotic phyla in the gene distribution search, and so eukaryotic genes were excluded from this analysis. The number of hits for each gene can be found in **Table 2**. These genes were aligned using MUSCLE v3.8.31 (*79*) then trimmed using TrimAl v.1.3 (*80*) with the -automated1 option as implemented in Phylemon2 (*91*). The model of evolution was determined by the Model Selection tool implemented in IQ-TREE 2.0.3 (*92*) with the flags *-m MFP, -mrate E,I,G,I+G,R and -madd C10,C20,C30,C40,C50,C60,EX2,EX3,EHO,UL2,UL3,EX_EHO,LG4M,LG4X,CF4,LG+C10,LG+ C20,LG+C30,LG+C40,LG+C50,LG+C60* to include all complex mixture models of protein evolution in model selection. All trees were constructed using IQ-TREE 2.0.3 (*92*) with a specification of 1000 ultrafast bootstraps (except aprB, which was set to 2000) and the default UFBoot convergence criterion. Within IQ-TREE, the “UFBoot stopping rule” automatically assesses the convergence of the split support values and stops collecting candidate trees once convergence is achieved (*93*). All gene trees reached convergence. The number of bootstraps used for each gene tree is reported in **Table 2**.

**Table 2.**
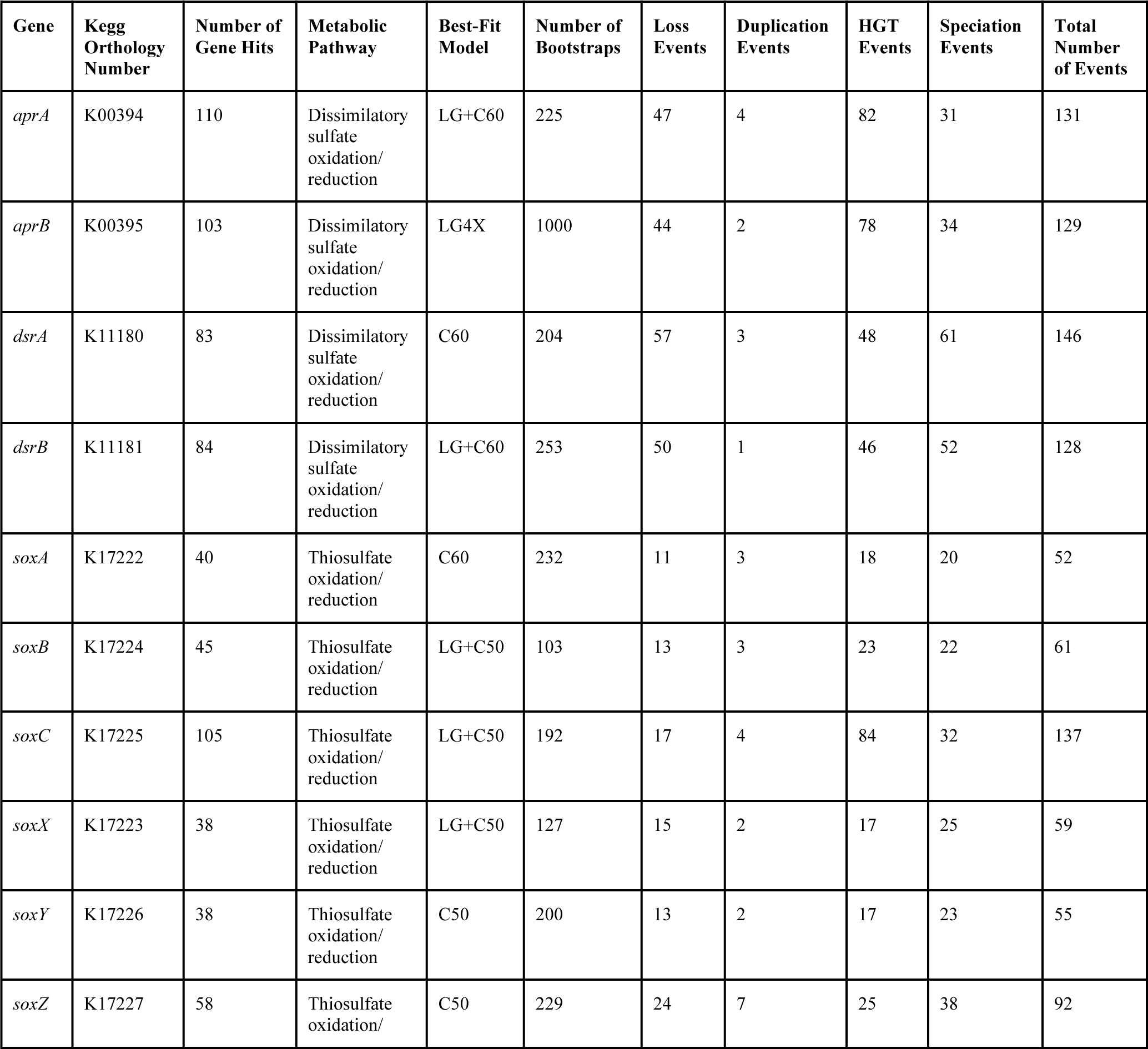
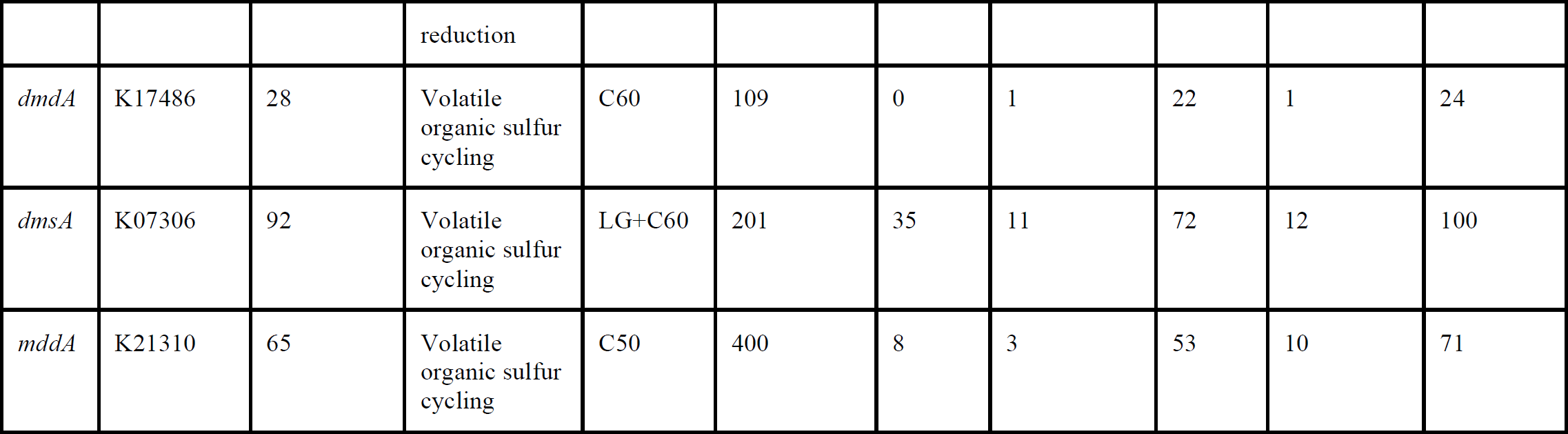
Sulfur cycling genes analyzed in this study. Number of gene hits indicates the number of genes identified among the genomes included in the species tree; Best-fit Model indicates the model of evolution used for generating the gene tree as determined by IQ-Tree ModelFinder; number of bootstraps indicates the number of bootstrap trees created to achieve convergence which were subsequently used for reconciliation in ecceTERA; number of speciation, loss, duplication, and HGT events are reported as determined by ecceTERA using the CIR clock model.

### Reconciliation of gene trees with species chronogram with ecceTERA

Gene trees and species trees were reconciled using ecceTERA v1.2.5 (*47*) to identify gene loss, duplication, speciation, and transfer events. We used the default settings implemented in ecceTERA and amalgamated the gene trees (amalgamate=true). No transfers to the dead were allowed when reconciling gene and species trees. The output was configured to recPhyloXML format (*94*) with the option “recPhyloXML.reconciliation=true”. Reconciliation analyses were performed on fully dated species trees and full sets of gene tree bootstraps (see **Table 3** for bootstrap information). Using a combination of custom Python scripts developed for this project (provided on GitHub at https://github.com/carleton-spacehogs/sulfur as well as on FigShare at 10.6084/m9.figshare.23255627), we calculated the mean date for each event based on the midpoints of the two 95% confidence intervals that defined the nodes of the branch on which the event occurred. Distributions of the gene event data produced from ecceTERA were subsequently compared to distribution data of the gene event data produced from AnGST (*46*) to ensure the results were not dependent on the reconciliation algorithm; the overall trends we observed were the same.

**Table 3.**
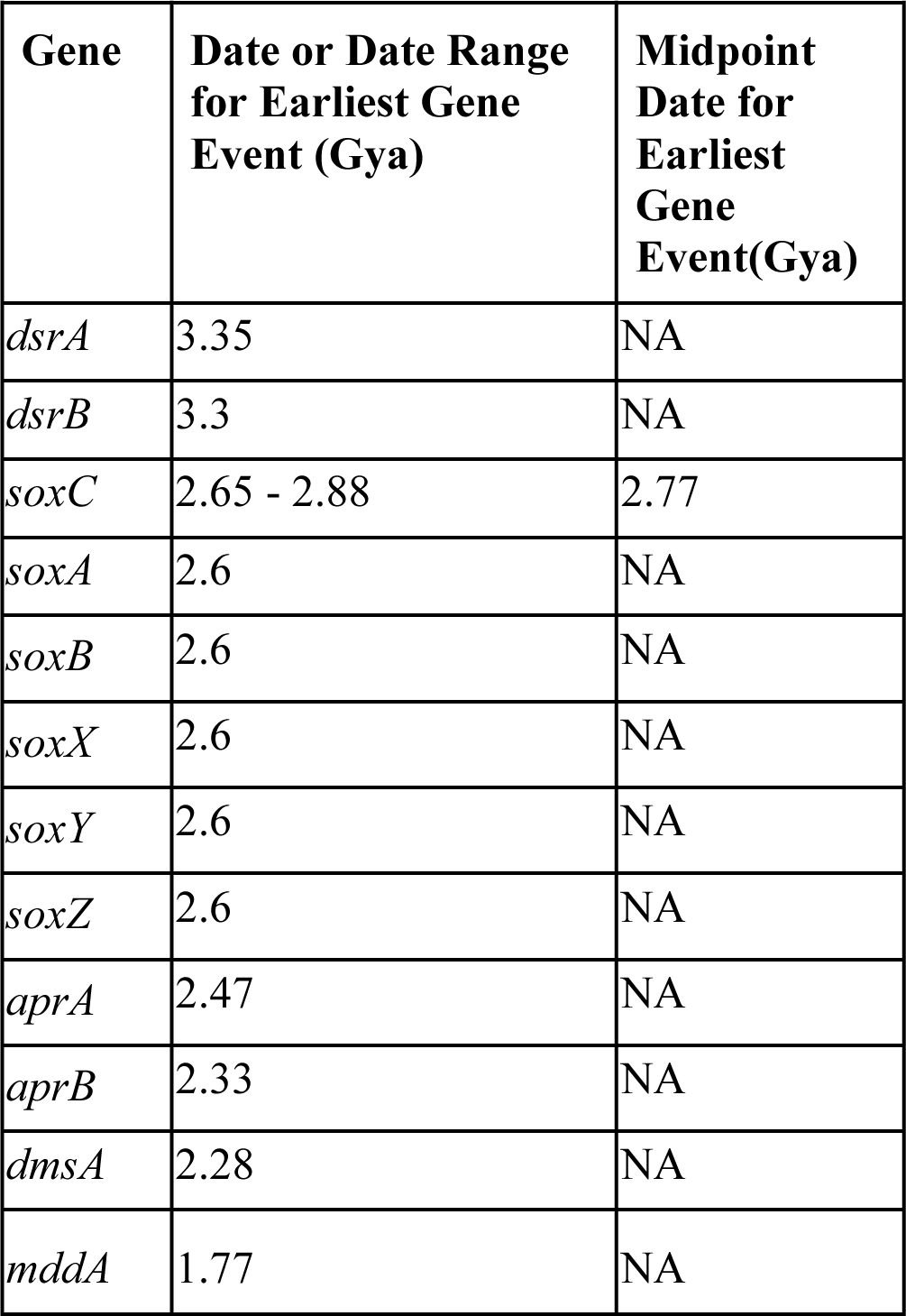
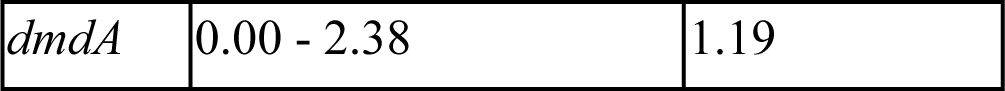
Identification of the earliest date for gene loss, duplication, speciation, and horizontal gene transfer events according to reconciliation with the chronogram generated using the CIR clock model and conservative calibration points, as identified by ecceTERA. While gene births are not inferred using this reconciliation method, the gene must have emerged by this point in time. Speciation events can be located to a precise node and therefore have a specific date, whereas duplication, loss, and transfer events occur on branches and are reported with a date range. For those events, we report the “midpoint” date between these two node dates in the right-hand column. NA, not applicable because these are speciation events at single nodes.

| **Gene** | **Date or Date Range for Earliest Gene Event (Gya)** | **Midpoint Date for Earliest Gene Event(Gya)** |
| --- | --- | --- |
| *dsrA* | 3.35 | NA |
| *dsrB* | 3.3 | NA |
| *soxC* | 2.65 - 2.88 | 2.77 |
| *soxA* | 2.6 | NA |
| *soxB* | 2.6 | NA |
| *soxX* | 2.6 | NA |
| *soxY* | 2.6 | NA |
| *soxZ* | 2.6 | NA |
| *aprA* | 2.47 | NA |
| *aprB* | 2.33 | NA |
| *dmsA* | 2.28 | NA |
| *mddA* | 1.77 | NA |
| *dmdA* | 0.00 - 2.38 | 1.19 |

## Acknowledgements

We thank Celine Scornavacca for assistance with ecceTERA, and Karthik Anantharaman for advice regarding the biological sulfur cycle.

## Author Contributions

RA and ES conceptualized the study. KM, GC, AK, BL, JS, and RA performed bioinformatics and phylogenetics analyses. KM, ES and RA wrote the manuscript with input from all authors. ES and RA supervised. ES and RA provided funding.

## Data and Materials Availability

All data needed to evaluate the conclusions in the paper are present in the paper and/or the Supplementary Materials.

