## Supplementary Materials for "The evolution and spread of sulfur-cycling enzymes reflect the redox state of the early Earth"

**Supplementary Table 1**. Identification of gene loss, duplication, and horizontal gene transfer events for all three clock models (UGAM, LN and CIR) using conservative calibration points, as determined by ecceTERA.

| | UGAM Clock Model | | | | | LN Clock Model | | | | |
|---|---|---|---|---|---|---|---|---|---|---|
| **Gene** | **Loss** | **Duplication** | **HGT** | **Speciation** | **Total** | **Loss** | **Duplication** | **HGT** | **Speciation** | **Total** |
| *aprA* | 25 | 4 | 77 | 46 | 152 | 21 | 4 | 80 | 40 | 145 |
| *aprB* | 9 | 2 | 80 | 26 | 117 | 21 | 2 | 72 | 42 | 137 |
| *dsrA* | 22 | 3 | 50 | 45 | 120 | 32 | 3 | 47 | 58 | 140 |
| *dsrB* | 27 | 2 | 50 | 49 | 128 | 31 | 2 | 48 | 54 | 135 |
| *soxA* | 11 | 3 | 18 | 20 | 52 | 11 | 3 | 18 | 20 | 52 |
| *soxB* | 13 | 3 | 24 | 22 | 62 | 13 | 3 | 24 | 22 | 62 |
| *soxC* | 26 | 4 | 81 | 36 | 141 | 26 | 4 | 81 | 36 | 141 |
| *soxX* | 15 | 2 | 16 | 25 | 58 | 15 | 2 | 16 | 25 | 58 |
| *soxY* | 13 | 2 | 17 | 23 | 55 | 13 | 2 | 17 | 23 | 55 |
| *soxZ* | 24 | 7 | 25 | 38 | 91 | 24 | 7 | 25 | 38 | 92 |
| *dmdA* | 0 | 1 | 22 | 1 | 24 | 0 | 1 | 22 | 1 | 24 |
| *dmsA* | 2 | 11 | 72 | 8 | 93 | 7 | 10 | 72 | 15 | 104 |
| *mddA* | 5 | 3 | 51 | 10 | 69 | 5 | 3 | 57 | 10 | 75 |

**Supplementary Table 2.** Identification of the earliest date for gene loss, duplication, speciation, and horizontal gene transfer events according to reconciliation with the chronogram generated using the LN and UGAM clock models and conservative calibration points, as identified by ecceTERA. While gene births are not inferred using this reconciliation method, the gene must have emerged by this point in time. Speciation events can be located to a precise node and therefore have a specific date, whereas duplication, loss, and transfer events occur on branches and are reported with a date range. For those events, we report the "midpoint" date between these two node dates in the right-hand column. NA, not applicable because these are speciation events at single nodes.

| | ***LN Clock Model*** | | ***UGAM Clock Model*** | |
|---|---|---|---|---|
| **Gene** | **Date or Date Range for Earliest Event (Gya)** | **Midpoint Date for Earliest Event (Gya)** | **Date Range for Earliest Event (Gya)** | **Midpoint Date for Earliest Event (Gya)** |
| aprA | 3.31 | NA | 1.89 - 2.60 | 2.24 |
| aprB | 3.46 | NA | 2.00 | NA |
| dsrA | 3.65 | NA | 2.94 | NA |
| dsrB | 3.24 - 3.43 | 3.33 | 2.38 - 2.62 | 2.50 |
| soxA | 2.76 | NA | 2.74 | NA |
| soxB | 2.76 | NA | 2.74 | NA |
| soxC | 2.96 - 3.18 | 3.07 | 2.25 - 2.50 | 2.38 |
| soxX | 2.76 | NA | 2.74 | NA |
| soxY | 2.76 | NA | 2.74 | NA |
| soxZ | 2.76 | NA | 2.74 | NA |
| dmdA | 0.0 - 3.09 | 1.54 | 0.00 - 2.12 | 1.06 |
| dmsA | 2.94 | NA | 1.97 | NA |
| mddA | 2.53 | NA | 0.0 - 3.03 | 1.51 |

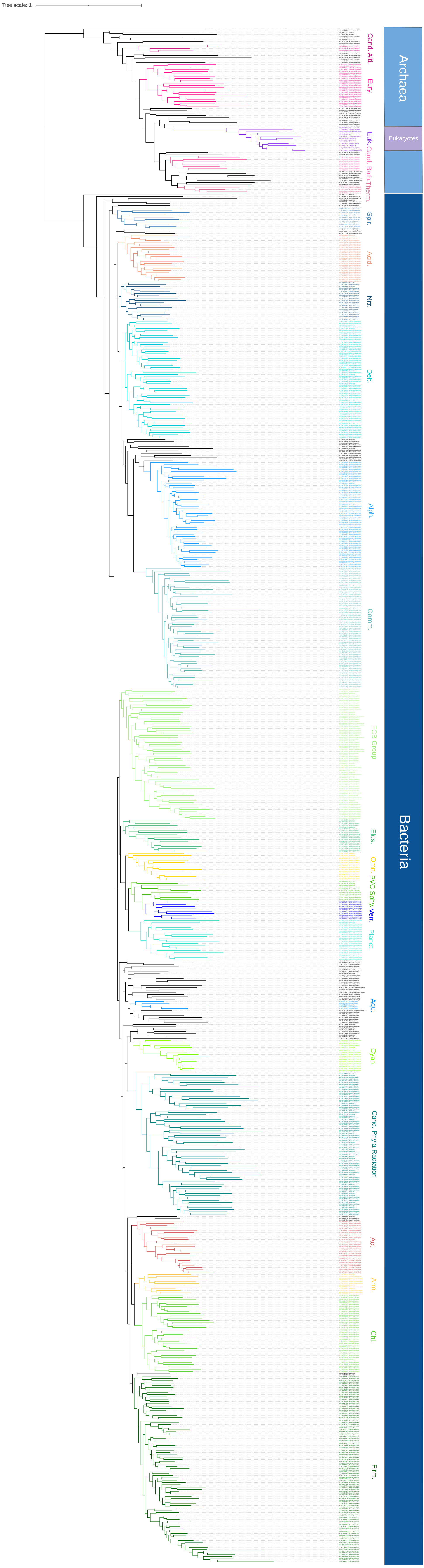

Tree scale: 1
Cand. Asli.
Eury.
Euk.
Cand. Bath.
Therm.
Spir.
Acid.
Nitr.
Delt.
Alph.
Gamm.
FCB Group
Elus.
Omn.
PVC
Sphy.
Verr.
Planct.
Aqu.
Cyan.
Cand. Phyla Radiation
Act.
Arm.
Chl.
Firm.
Archaea
Eukaryotes
Bacteria

**Supplementary Figure 1.** Tree of life ("species tree") used for this study. The tree includes 871 genomes in total, including 777 bacterial, 80 archaeal, and 14 eukaryotic genomes. The bacterial and archaeal genomes represent one genome per order based on the GTDB taxonomy.

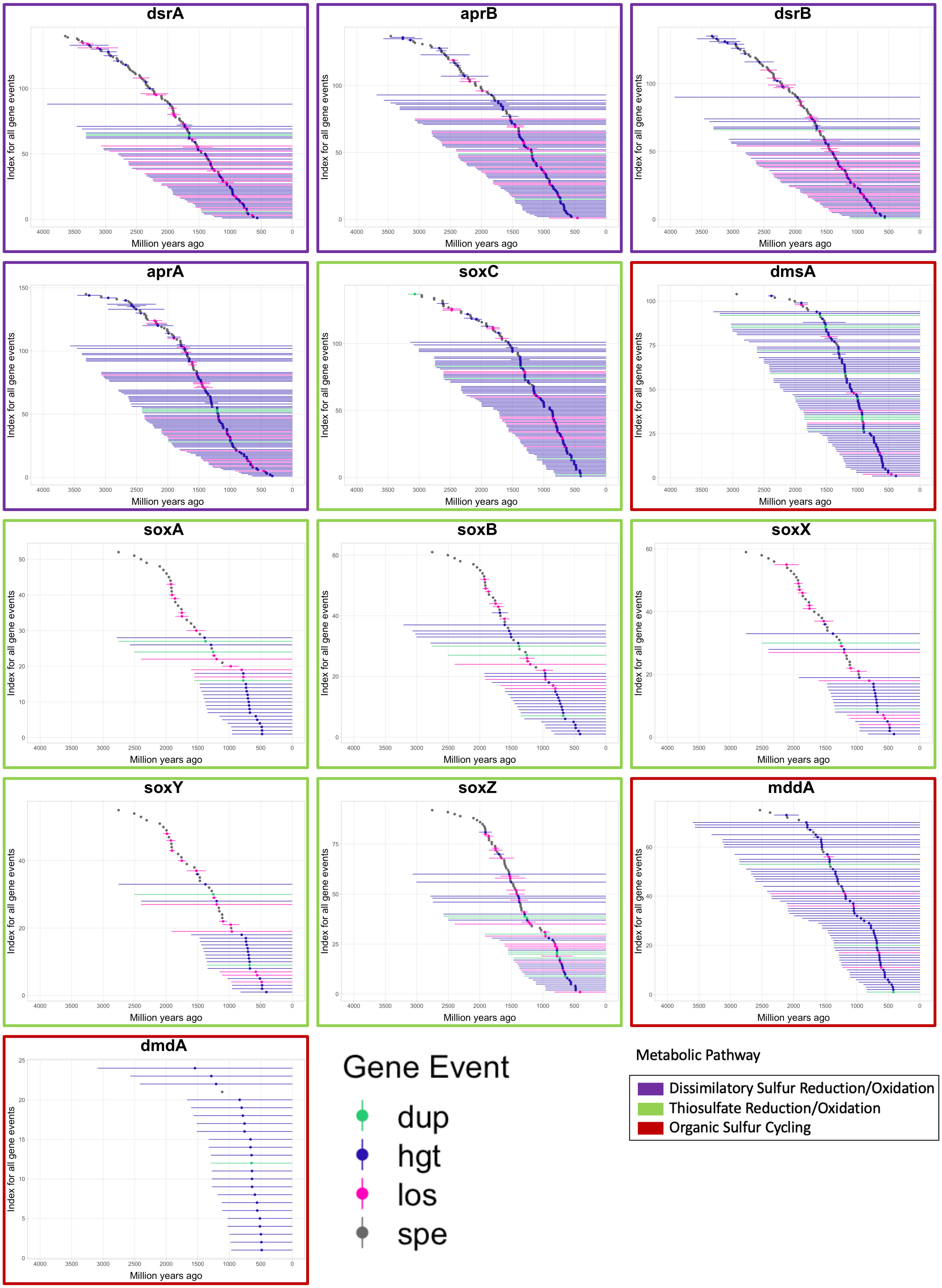

dsrA
aprB
dsrB
aprA
soxC
dmsA
soxA
soxB
soxX
soxY
soxZ
mddA
dmdA
Index for all gene events
Million years ago
Gene Event
dup
hgt
los
spe
Metabolic Pathway
Dissimilatory Sulfur Reduction/Oxidation
Thiosulfate Reduction/Oxidation
Organic Sulfur Cycling

**Supplementary Figure 2**. Time ranges for each gene duplication, transfer, loss and speciation event identified for all sulfur-cycling genes. Reconciliations were conducted with the lognormal clock model. The time ranges reflect the branch lengths for branches on the time-calibrated tree of life on which these events were identified; the event could have occurred anywhere along the branch. Graphs are placed in chronological order according to the midpoint of the earliest event. Dup =duplication, hgt = horizontal gene transfer, los = loss, spe = speciation event.

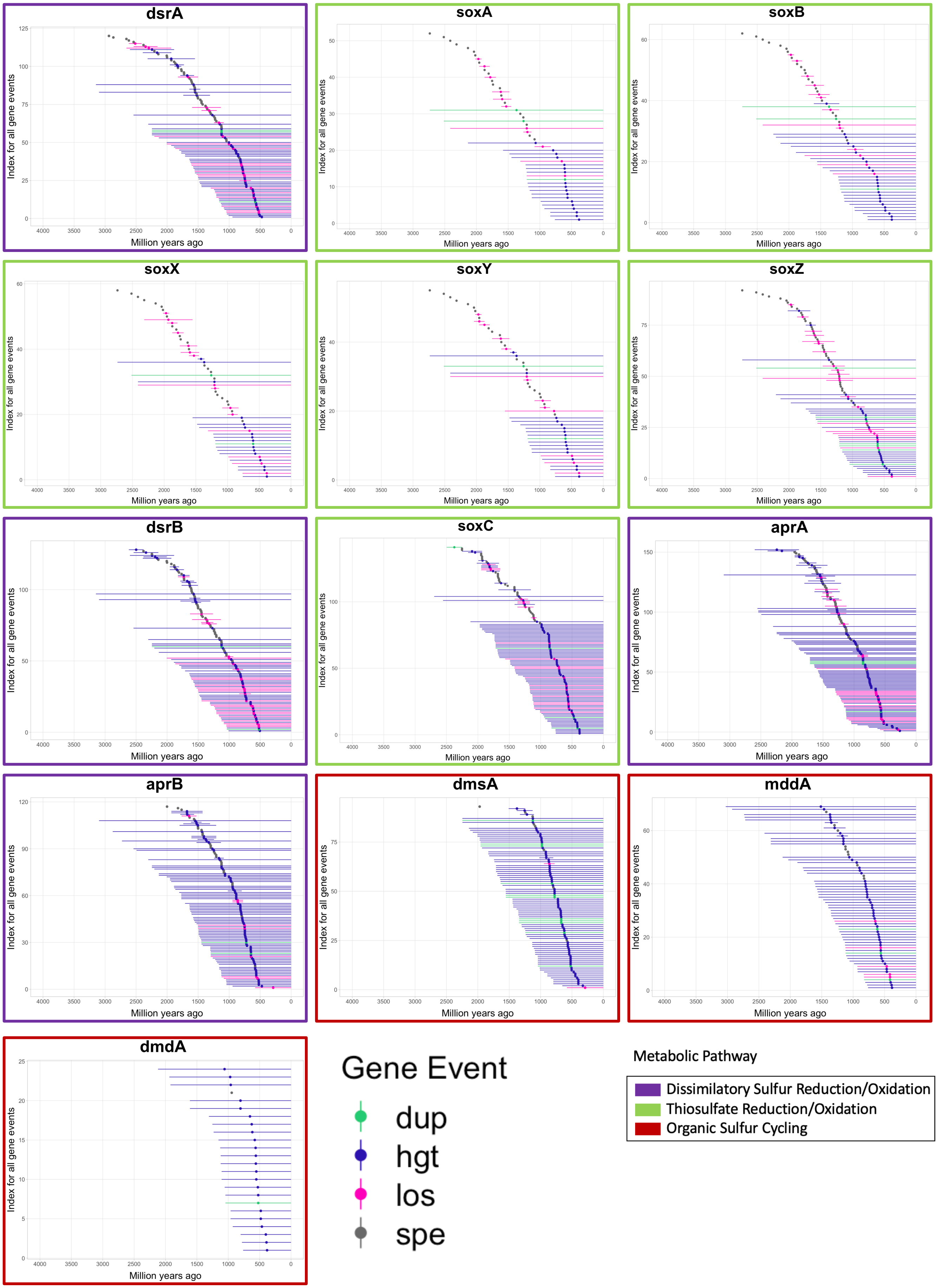

dsrA
soxA
soxB
soxX
soxY
soxZ
dsrB
soxC
aprA
aprB
dmsA
mddA
dmdA
Index for all gene events
Million years ago
4000 3500 3000 2500 2000 1500 1000 500 0
Gene Event
dup
hgt
los
spe
Metabolic Pathway
Dissimilatory Sulfur Reduction/Oxidation
Thiosulfate Reduction/Oxidation
Organic Sulfur Cycling

**Supplementary Figure 3**. Time ranges for each gene duplication, transfer, loss, and speciation event identified for all sulfur-cycling genes. Reconciliations were conducted with the UGAM clock model. The time ranges reflect the branch lengths for branches on the time-calibrated tree of life on which these events were identified; the event could have occurred anywhere along the branch. Graphs are placed in chronological order according to the midpoint of the earliest event. Dup =duplication, hgt = horizontal gene transfer, los = loss, spe = speciation event.

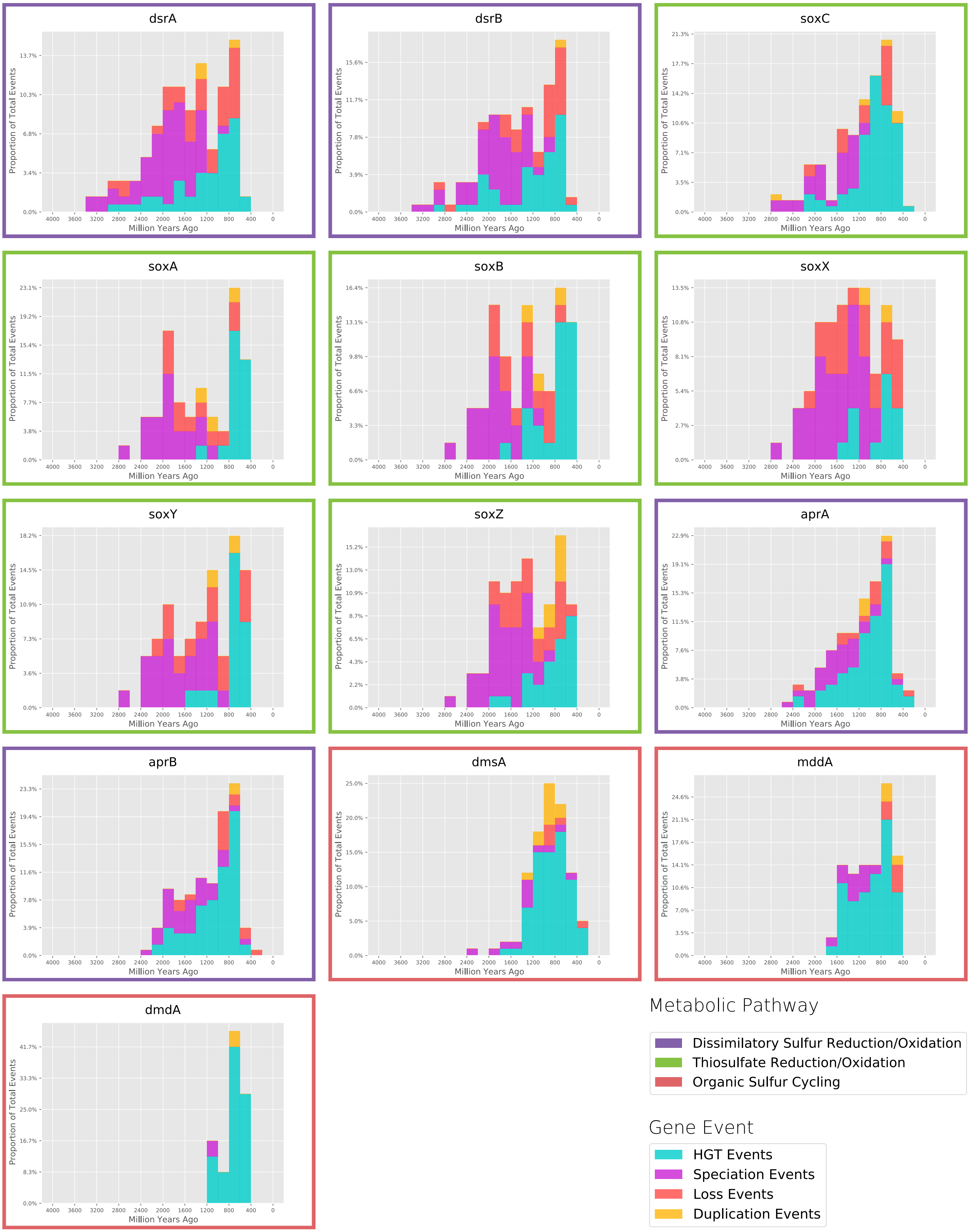
dsrA
dsrB
soxC
soxA
soxB
soxX
soxY
soxZ
aprA
aprB
dmsA
mddA
dmdA
Proportion of Total Events
Million Years Ago
Metabolic Pathway
Dissimilatory Sulfur Reduction/Oxidation
Thiosulfate Reduction/Oxidation
Organic Sulfur Cycling
Gene Event
HGT Events
Speciation Events
Loss Events
Duplication Events

**Supplementary Figure 4**. Histograms of gene loss, duplication, speciation, and HGT, as identified by ecceTERA using the CIR clock model. The y-axis represents the proportion of total gene events. Histogram bins are 200 million years in width. Colored boxes outlining each graph represent the general metabolic pathway in which each gene belongs. Histograms are placed in chronological order according to the midpoint of the earliest event.

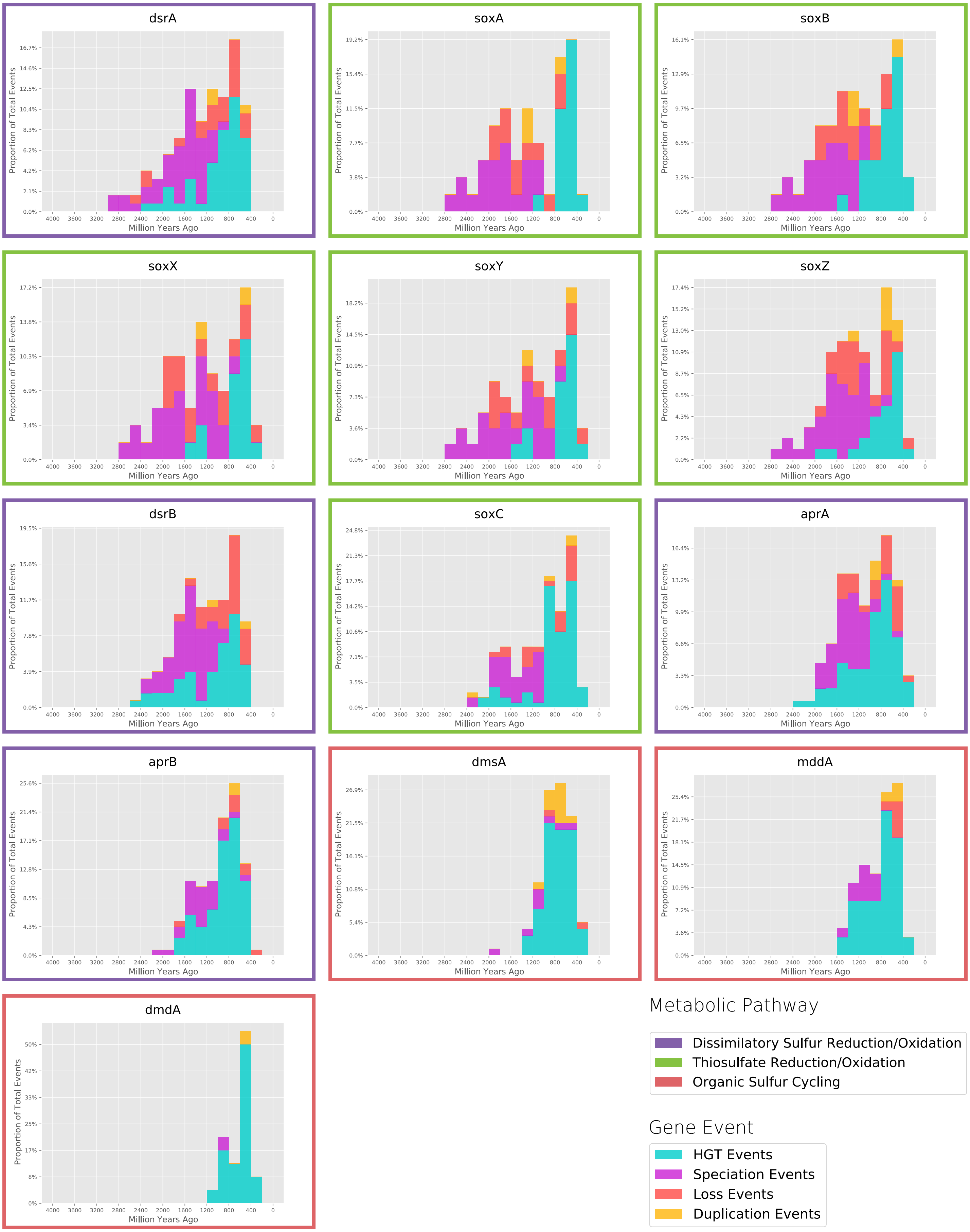
dsrA
soxA
soxB
soxX
soxY
soxZ
dsrB
soxC
aprA
aprB
dmsA
mddA
dmdA
Proportion of Total Events
Million Years Ago
4000 3600 3200 2800 2400 2000 1600 1200 800 400 0
Metabolic Pathway
Dissimilatory Sulfur Reduction/Oxidation
Thiosulfate Reduction/Oxidation
Organic Sulfur Cycling
Gene Event
HGT Events
Speciation Events
Loss Events
Duplication Events

**Supplementary Figure 5.** Histograms of gene loss, duplication, speciation, and HGT, as identified by ecceTERA using the UGAM clock model. The y-axis represents the proportion of total gene events. Histogram bins are 200 million years in width. Colored boxes outlining each graph represent the general metabolic pathway in which each gene belongs. Histograms are placed in chronological order according to the midpoint of the earliest event.

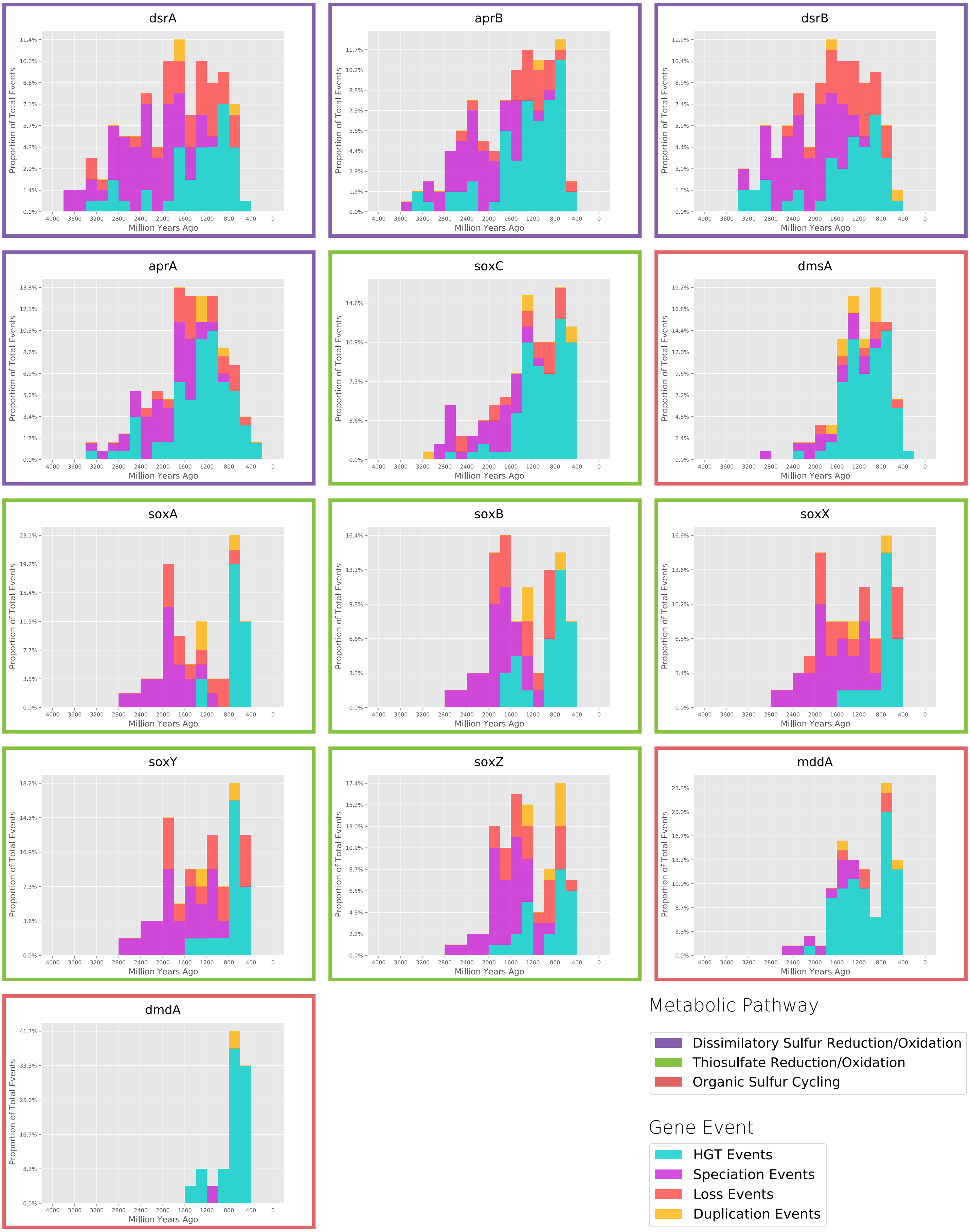
dsrA
aprB
dsrB
aprA
soxC
dmsA
soxA
soxB
soxX
soxY
soxZ
mddA
dmdA
Proportion of Total Events
Million Years Ago
4000 3600 3200 2800 2400 2000 1600 1200 800 400 0
Metabolic Pathway
Dissimilatory Sulfur Reduction/Oxidation
Thiosulfate Reduction/Oxidation
Organic Sulfur Cycling
Gene Event
HGT Events
Speciation Events
Loss Events
Duplication Events

**Supplementary Figure 6**. Histograms of gene loss, duplication, speciation, and HGT, as identified by ecceTERA using the lognormal clock model. The y-axis represents the proportion of total gene events. Histogram bins are 200 million years in width. Colored boxes outlining each graph represent the general metabolic pathway in which each gene belongs. Histograms are placed in chronological order according to the midpoint of the earliest event.
